# Hippocampo–supramammillary coupling across sleep and wake

**DOI:** 10.1101/2025.10.08.681248

**Authors:** M. Caneo, N. Espinosa, G. Lazcano, A. Aguilera, A. Lara-Vasquez, P. Fuentealba

**Affiliations:** Laboratory of Neural Circuits, Departamento de Psiquiatria, Facultad de Medicina, Pontificia Universidad Catolica de Chile

## Abstract

Hippocampal communication with subcortical circuits reorganizes across sleep–wake states, yet the dynamics of such interactions remain incompletely defined. We simultaneously recorded neuronal spiking and local field potentials from the dorsal hippocampus (CA1 and dentate gyrus) and limbic subcortical nuclei (supramammillary nucleus (SuM) and lateral septum (LS)) in freely behaving rats across the sleep–wake cycle. During quiescent states (non-rapid eye movement (nREM) sleep and quiet wakefulness), sharp-wave ripples dominated hippocampal activity and produced fast, top–down excitation, with large population surges locally in CA1 and DG, and more modest but significant activation in LS and SuM, whereas dentate spikes elicited smaller state-invariant responses. Conversely, in the bottom–up direction, epochs of high-discharge SuM activity were associated with slower, state-dependent activation of hippocampal populations that was larger in wake than in sleep, revealing distinct temporal scales and state dependence for reciprocal pathways. During activated, theta-enriched states (rapid eye movement (REM) sleep and active wakefulness), spike–field coupling revealed circuit-wide theta coordination. In active wakefulness, SuM bursts produced brief inhibition of CA1 spiking while theta oscillations organized a multiregional firing sequence around the theta cycle trough; yet SuM neurons were not significantly phase-locked. In REM sleep, this pattern inverted, with only SuM neurons significantly phase-locking to theta waves, and preferred firing near the theta cycle peak. Together, these findings identify state-dependent, bidirectional coordination between hippocampus and subcortical nuclei, characterized by ripple-locked top–down hippocampal output during quiescent states and SuM-mediated bottom–up modulation that reconfigures under theta during activated states.

## INTRODUCTION

Hippocampal sharp-wave ripples (100–250 Hz) are among the most synchronous population events in the mammalian brain (1), prominent during non-rapid eye movement (nREM) sleep and quiet wakefulness (2, 3), and widely implicated in systems-level memory consolidation and the routing of recent experience during wake (4). Converging evidence indicates that during hippocampal ripples, coordinated thalamocortical activity establishes a protected window for hippocampal–cortical communication by transiently suppressing output from subcortical regions involved in sensory processing or procedural learning (5). This gating is thought to minimize interference and thereby facilitate consolidation of hippocampus-dependent memories; consistent with this view, broad regions of the cerebral cortex are selectively activated during ripple events, whereas many diencephalic, midbrain, and brainstem nuclei are strongly and consistently inhibited (5, 6). Causal disruption studies and recent syntheses converge on a necessary role for ripples in memory (7, 8), and ripple features covary with slower, brain-wide excitability fluctuations that predict diverse extra-hippocampal responses (6, 9). In addition to ripples, dentate spikes are prominent dentate gyrus (DG) fast events thought to reflect synchronous perforant-path input (10); dentate spikes reorganize excitation–inhibition across the trisynaptic circuit and are increasingly implicated in offline processing. Recent studies classify dentate spikes subtypes, demonstrate dentate spike-linked reactivation and broad network coupling, and suggest mnemonic roles that complement ripple-supported replay (11–13), although the downstream impact and state dependence of dentate spikes remain less well established than for ripple episodes.

Two subcortical limbic nodes intimately coupled to the hippocampus are the supramammillary nucleus (SuM) and the lateral septum (LS). SuM sends dense projections to DG, CA1, and CA2, with collaterals reaching the septal complex (14–16). These pathways can pace or modulate hippocampal theta oscillations and influence arousal and REM sleep expression (17). Recent circuit studies reveal mixed glutamatergic and GABAergic transmission from SuM, the DG projection controls theta/gamma power and synaptic plasticity, the CA2 projection contributes to social memory, while the functional implications of the recently identified CA1 direct projection remain to be described (14–16). During behavior, SuM activity covaries with hippocampal dynamics, as SuM-evoked DG calcium signals become tightly correlated during spatial memory retrieval (18). On the other hand, long considered as a relay, LS is now recognized as a multifunctional hub with rich, topographically organized connectivity linking hippocampus to hypothalamic and midbrain systems that regulate motivation, affect, and goal-directed behaviors (19–21). Anatomical mapping show precise hippocampo–septal topography, and recent work details parallel hippocampo–septal pathways that differentially bias approach–avoidance, as well as LS influences on dopaminergic circuits (22, 23). Functionally, LS carries spatial and locomotor variables and integrates cognitive with internal-state signals across behaviors (20).

Here we combined simultaneous multisite recordings of neuronal spiking and local field potentials from the dorsal hippocampus (CA1 and DG) and subcortical nuclei (SuM and LS) in freely behaving rats to examine their coordination across the sleep–wake cycle. During quiescent states (nREM sleep and quiet wake (24)), sharp-wave ripples drove large excitatory surges in CA1 and DG with weaker co-activation of LS and SuM in the top–down direction, whereas dentate spikes elicited smaller, largely state-invariant responses. In the bottom–up direction, high-discharge SuM epochs predicted activation of hippocampal populations that was stronger in wake than in sleep. During activated states (REM sleep and active wake (25)), hippocampal theta oscillations coordinated the limbic circuit, with SuM bursts producing rapid inhibition of CA1, but not DG, selectively during wake, and SuM neurons phase-locked to theta specifically in REM, but not during active wake. Together, these findings show that hippocampal–subcortical coordination is bidirectional and state-dependent, combining fast, ripple-locked top–down output with rapid, SuM-mediated bottom–up modulation that reconfigures under hippocampal theta oscillations.

## RESULTS

### Hippocampal ripples coordinate state-invariant spiking responses in the supramammillary nucleus

To characterize functional interactions between the dorsal hippocampus and subcortical regions (**Table S1, Fig. S1**), we simultaneously recorded neuronal activity in dorsal hippocampus (CA1 and DG) and subcortical associated nuclei (SuM and LS) in freely behaving rats (**Fig. 1A**). Recordings spanned spontaneous alternations across the sleep–wake cycle (**Table S2**). Brain-states were identified from hypnograms that combined the hippocampal power spectrum with movement speed derived from video tracking (**Fig. 1B**). We also overlaid locomotor speed and SuM population firing rates (**Fig. 1B**), which were significantly correlated (**Fig. 1C**). Across regions, firing-rate distributions were broadly similar across brain-states, only the SuM showed clear state dependence, with lower firing rates during nREM sleep (**Fig. 1D**). We next extracted hippocampal sharp-wave ripples and used the onsets to assess temporal coordination between hippocampus and subcortical targets (**Fig. 1A**). Ripple-triggered time–frequency maps revealed in each recorded region, a brief broadband power increase dominated by a narrow, vertical enhancement in the high-frequency range (>100 Hz), suggestive of spike-locked transients to hippocampal ripples (**Fig. S2**). In DG, high-frequency power concentrations were also consistent with the presence of dentate spikes (**Fig. S2**). In SuM, ripple onset was accompanied by a prominent increase in low-gamma activity (20–40 Hz, **Fig. S2**) which reflected a phase-locked evoked potential (**Fig. 1E**). Such event related potential, expressed as a negative deflection in the local field potential, was present in all recorded animals to a different extent (**Fig. 1E**) and varied in magnitude with brain-state, being smaller during quiet wakefulness than during nREM sleep (Wilcoxon signed-rank, p = 0.009, **Fig. 1E**). However, peak latencies remain consistent regardless of brain-state (Wilcoxon signed-rank, p = 0.297, **Fig. 1E**). Together, these observations point to a significant, state-dependent synaptic drive from dorsal hippocampus to SuM during ripple episodes.

**Figure 1.**
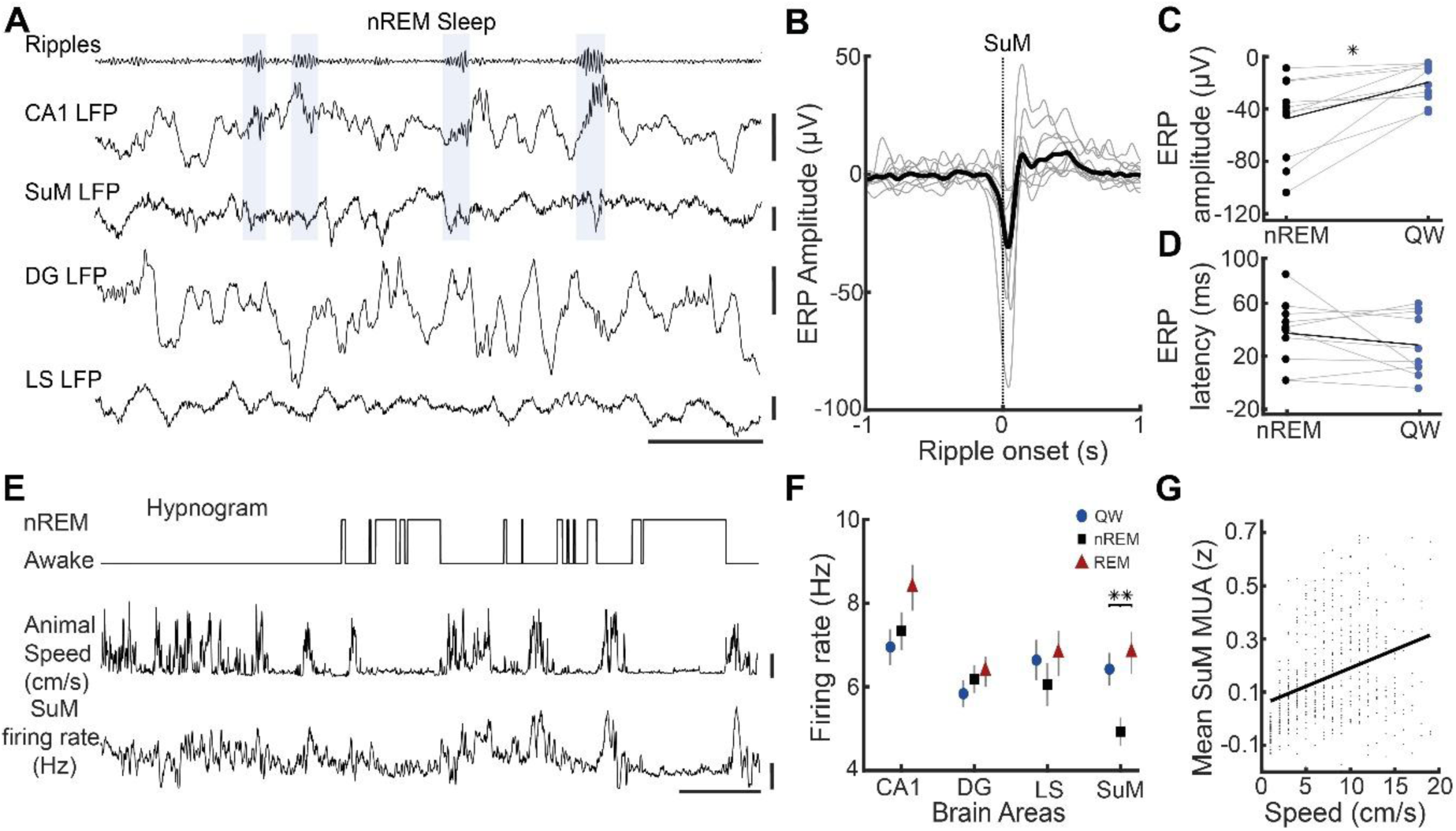
State-dependent hippocampo–subcortical coupling and SuM activity. **A,** Representative simultaneous LFPs from dorsal CA1 (CA1d), supramammillary nucleus (SuM), dentate gyrus (DG) and lateral septum (LS). Blue boxes mark CA1d ripple epochs; note concomitant deflections, particularly in SuM. Scale bars at right; same animal. **B,** Ripple-triggered average of SuM LFP aligned to CA1d ripple onset (t = 0) during nREM; thin gray traces, individual animals; thick line, population mean. **C–D,** Across animals, SuM ripple-locked field responses show larger amplitude in nREM than in quiet wake (QW) (paired comparison; *p* < 0.05) with no change in onset latency (*p* > 0.05). **E,** Example hypnogram (awake vs nREM) from a recording segment with the animal’s speed (cm/s) and simultaneously estimated SuM firing rate (Hz); SuM activity increases during waking/locomotion. **F,** Mean firing rates by brain region and vigilance state (QW, circles; nREM, squares; REM, triangles). Error bars, SEM. A state effect is evident in SuM (**, *p* < 0.01). **G,** Relationship between locomotor activity and SuM neuronal discharge: mean SuM multi-unit activity (MUA; z-score) versus running speed shows a positive correlation across time bins. QW, quiet wake; nREM, non-REM sleep; SuM, supramammillary nucleus; DG, dentate gyrus; LS, lateral septum.

We next studied the ripple-associated spiking across hippocampal and subcortical circuits (**Table S3**). Ripples robustly modulated both local (hippocampal) and distal (subcortical) firing, yielding a net excitatory effect across regions (**Fig. 2A**). At the single-unit level, responses were heterogeneous, yet nearly half of CA1 neurons were significantly excited by ripples (49.4%; **Fig. 2A**). In DG, a smaller yet substantial fraction of units was activated (39.1%; **Fig. 2A**). Subcortical nuclei were less strongly modulated than hippocampus but nonetheless showed population-level excitation, with over a quarter of SuM neurons (28.7%) and a smaller proportion in LS (19.6%) significantly activated (**Fig. 2A**). Population peri-stimulus time histograms revealed the largest ripple-locked surge in CA1, which was stronger during sleep than wake (peak amplitude: nREM, 3.62 ± 0.34 z; QW, 2.99 ± 0.29 z; Wilcoxon signed-rank test, FDR-corrected, p < 1.3x10^-7^; **Fig. 2B**). DG exhibited a weaker yet clear activation, with no significant brain-state differences (nREM, 1.11 ± 0.15 z; QW, 0.95 ± 0.15 z, Wilcoxon signed-rank test, FDR-corrected, p > 0.05; **Fig. 2B**). Consistent with circuit anatomy, ripple-evoked activation in DG lagged that in CA1 (peak latency, CA1, 40.8 ± 1.72 ms; DG, 54.3 ± 2.01 ms; Wilcoxon rank-sum test, p = 6.5x10^-7^). Among subcortical targets, SuM responses were comparable across brain-states (nREM, 0.95 ± 0.13 z, QW, 0.70 ± 0.15 z; Wilcoxon signed-rank test, FDR-corrected, p > 0.05; **Fig. 2B**), whereas LS displayed a brief, more modest increase that was likewise state-invariant (nREM, 0.45 ± 0.01 z; QW, 0.22 ± 0.09 z; Wilcoxon signed-rank test, FDR-corrected, p > 0.05; **Fig. 2B**).

**Figure 2.**
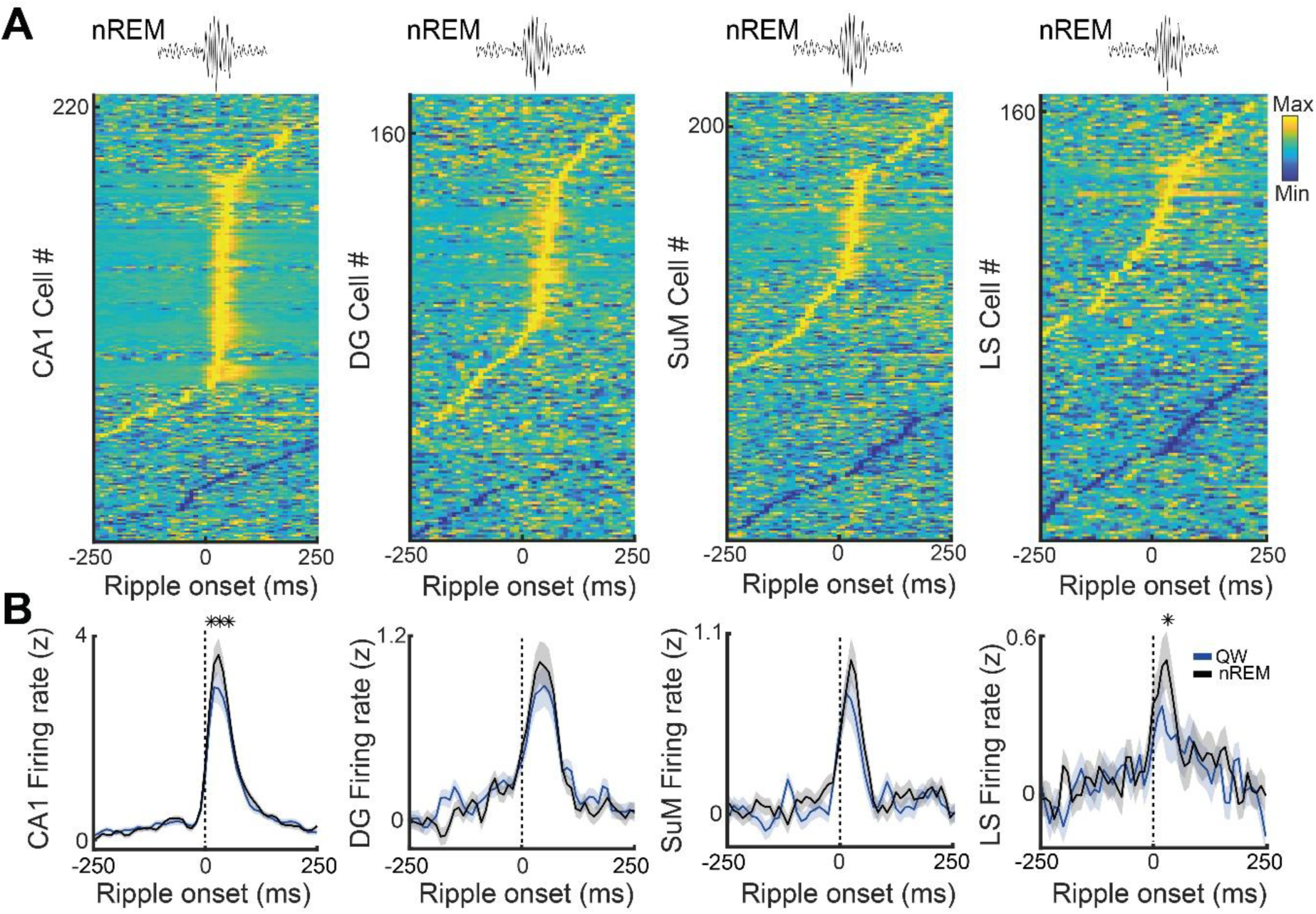
Ripple-locked spiking across hippocampal and subcortical populations during quiescent states. **A,** Heat maps of z-scored firing rates from single units in dorsal CA1 (CA1d), dentate gyrus (DG), supramammillary nucleus (SuM), and lateral septum (LS), each aligned to CA1 ripple onset (t = 0; vertical dashed line). One row = one unit; units are sorted by the latency of their peak modulation. Color scale indicates relative firing (Min→Max). Cartoon traces above panels show example ripple waveform used for alignment. Numbers on the y-axes indicate the number of units included per region. **B,** Population peri-stimulus time histograms (mean ± SEM; z-score) for quiet wake (QW, blue) and non-REM sleep (nREM, black). CA1 displays a sharp peak tightly time-locked to ripple onset, DG a moderate modulation, and SuM and LS weaker, delayed increases relative to CA1, consistent with attenuated subcortical recruitment during ripples. Ripple-locked spiking is broadly similar across QW and nREM (state-invariant) for DG and SuM, whereas CA1 and LS, show a small but significant increase. Asterisks denote significant differences between distributions (each * depicts FDR-corrected p < 0.05).

Dentate spikes from DG evoked weaker spiking that was largely state-invariant (**Fig. 3A**). Overall, DG dentate spikes elicited smaller population responses than CA1 ripple episodes (dentate spikes: 0.48 ± 0.04 z; ripples: 1.51 ± 0.09 z; F(1, 2593) = 86.1, p = 3.6x10^-20^, three-way ANOVA [brain state x brain region x hippocampal pattern]). In DG, dentate spike-triggered firing rose modestly with undetectable differences across brain-states (nREM, 0.74 ± 0.18 z; QW, 0.76 ± 0.14 z; Wilcoxon signed-rank test, FDR-corrected, p > 0.05; **Fig. 3B**). CA1 responses to dentate spikes were likewise modest and brain-state-invariant (nREM, 0.76 ± 0.10 z; QW, 0.56 ± 0.09 z, Wilcoxon signed-rank test, FDR-corrected, p > 0.05; **Fig. 3B**). As expected from synaptic connectivity, dentate spike activation in DG was faster than in CA1 (peak latency, CA1d, 40.0 ± 2.07 ms; DGd, 28.2 ± 2.21 ms; Wilcoxon rank-sum test, p = 6.1x10^-5^). Among subcortical targets, LS showed no significant modulation by dentate spikes (Wilcoxon signed-rank test against zero, FDR-corrected, p > 0.05; **Fig. 3B**), whereas SuM exhibited a small but significant increase in firing that remained state-invariant (nREM, 0.31 ± 0.09 z; QW, 0.32 ± 0.08 z; Wilcoxon signed-rank test, FDR-corrected, p > 0.05; peak latency; **Fig. 3B**). Together, these results indicate that dentate spikes elicit comparatively weak spiking both locally within hippocampus and in target subcortical nuclei, contrasting with the robust ripple-locked activation, whose subcortical magnitude remained largely state-invariant. Thus, despite clear state–dependent variation in hippocampal synaptic drive to SuM, the net spiking output of SuM remained essentially constant across states.

**Figure 3.**
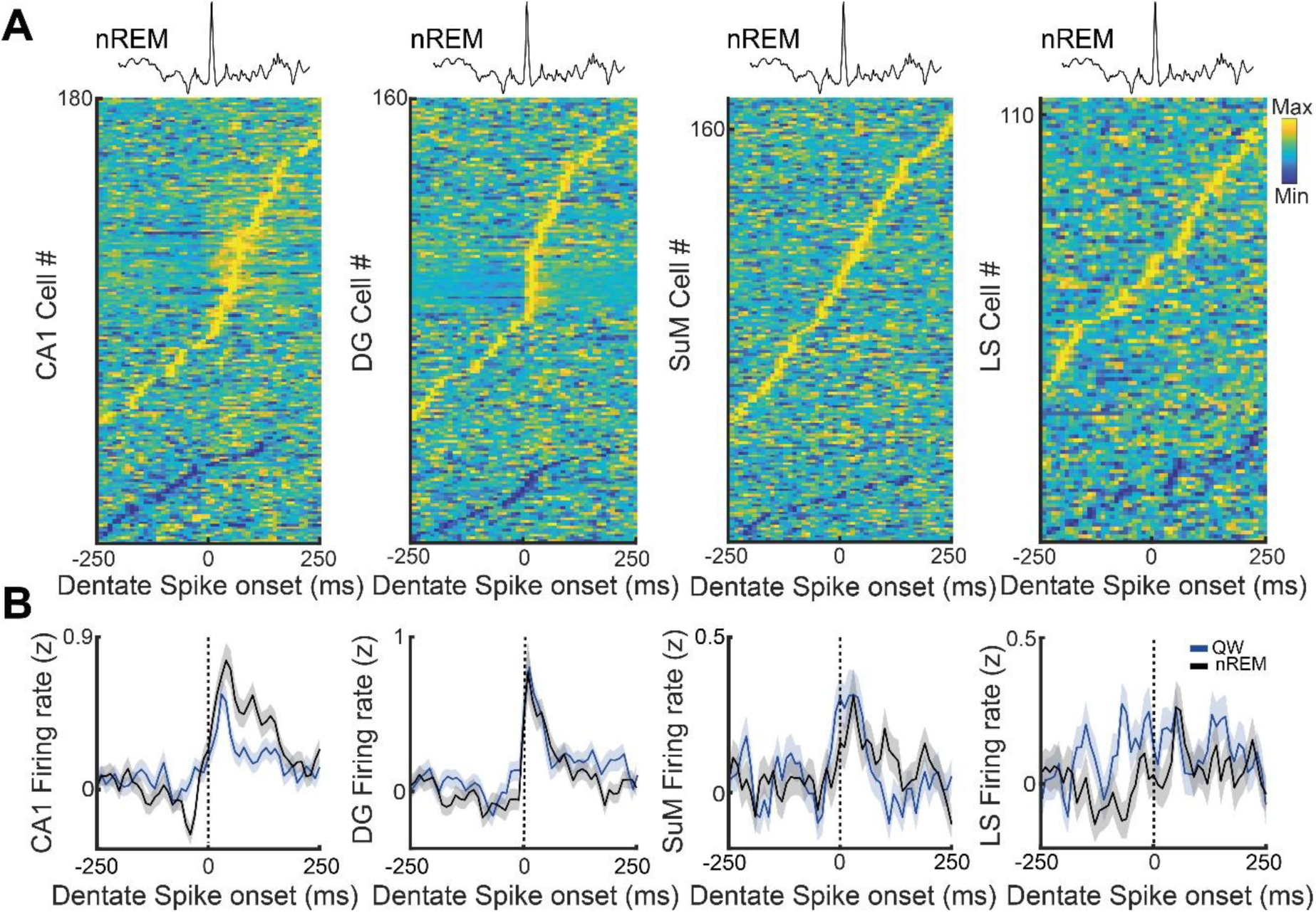
Dentate spike–locked spiking across hippocampal and subcortical populations during quiescent states. **A,** Heat maps of z-scored firing rates from single units in dorsal CA1 (CA1d), dentate gyrus (DG), supramammillary nucleus (SuM) and lateral septum (LS), aligned to dentate-spike (DS) onset in DG (t = 0; vertical dashed line). One row = one unit; units are sorted by the latency of peak modulation. Color scale indicates relative firing (Min→Max). Traces above panels show example DS waveform used for alignment. **B,** Population peri-stimulus time histograms (mean ± SEM; z-score) for quiet wake (QW, blue) and non-REM sleep (nREM, black). CA1 and DG exhibit significant, short-latency increases time-locked to DS, whereas subcortical regions show weaker (SuM) or no (LS) modulation. DS-evoked spiking is broadly similar across QW and nREM (state-invariant), consistent with a local effect that propagates only weakly to subcortical targets. Histogram distributions were not significantly different between brain-states (* would depict FDR-corrected p < 0.05).

### Supramammillary nucleus volleys drive dorsal hippocampus activity with state-dependent amplitude

To our knowledge, neither locally generated high-frequency oscillations (>100 Hz) nor ripple-like field potential events have been reported in SuM (26, 27). Accordingly, we used SuM multiunit activity (MUA) amplitude as a proxy for high-frequency bursting activity (**Fig. 4A**). Hence, we first calibrated a suprathreshold window by aligning SuM single-unit spikes to MUA excursions of increasing magnitude and determined the range at which local spiking activation saturated (**Fig. S3**). We then detected high-activity SuM MUA epochs within this range and used them to probe associated network effects. We found that when comparing the temporal profiles of population discharge across brain-states revealed a slower (seconds-scale) activation pattern throughout the circuit of larger amplitude during wake than during sleep. Within such state of generalized slow activation, peak firing rates locked to high-frequency SuM MUA epochs were state-invariant in SuM and LS (SuM: nREM, 3.4 ± 0.34 z; QW, 4.0 ± 0.36 z; Wilcoxon signed-rank test, FDR-corrected, p > 0.05; LS: nREM, 0.68 ± 0.12 z; QW, 0.55 ± 0.06 z; Wilcoxon signed-rank, FDR-corrected, p > 0.05; **Fig. 4B**). Conversely, aligning high-frequency SuM MUA to hippocampal activity revealed robust, state-dependent activation in CA1 and DG, with larger peak responses during wakefulness than during sleep (CA1: nREM, 0.47 ± 0.07 z; QW, 0.61 ± 0.06 z; Wilcoxon signed-rank test, FDR-corrected, p < 10⁻⁴; DG: nREM, 0.27 ± 0.07 z; QW, 0.50 ± 0.05 z; Wilcoxon signed-rank, FDR-corrected, p < 10⁻⁴; **Fig. 4B**). Together, these data indicate that brief elevations in SuM high-frequency discharge are associated with slower scale dynamics of increased overall circuit activity, and can elicit slow, brain-state–dependent population responses within hippocampal circuits.

**Figure 4.**
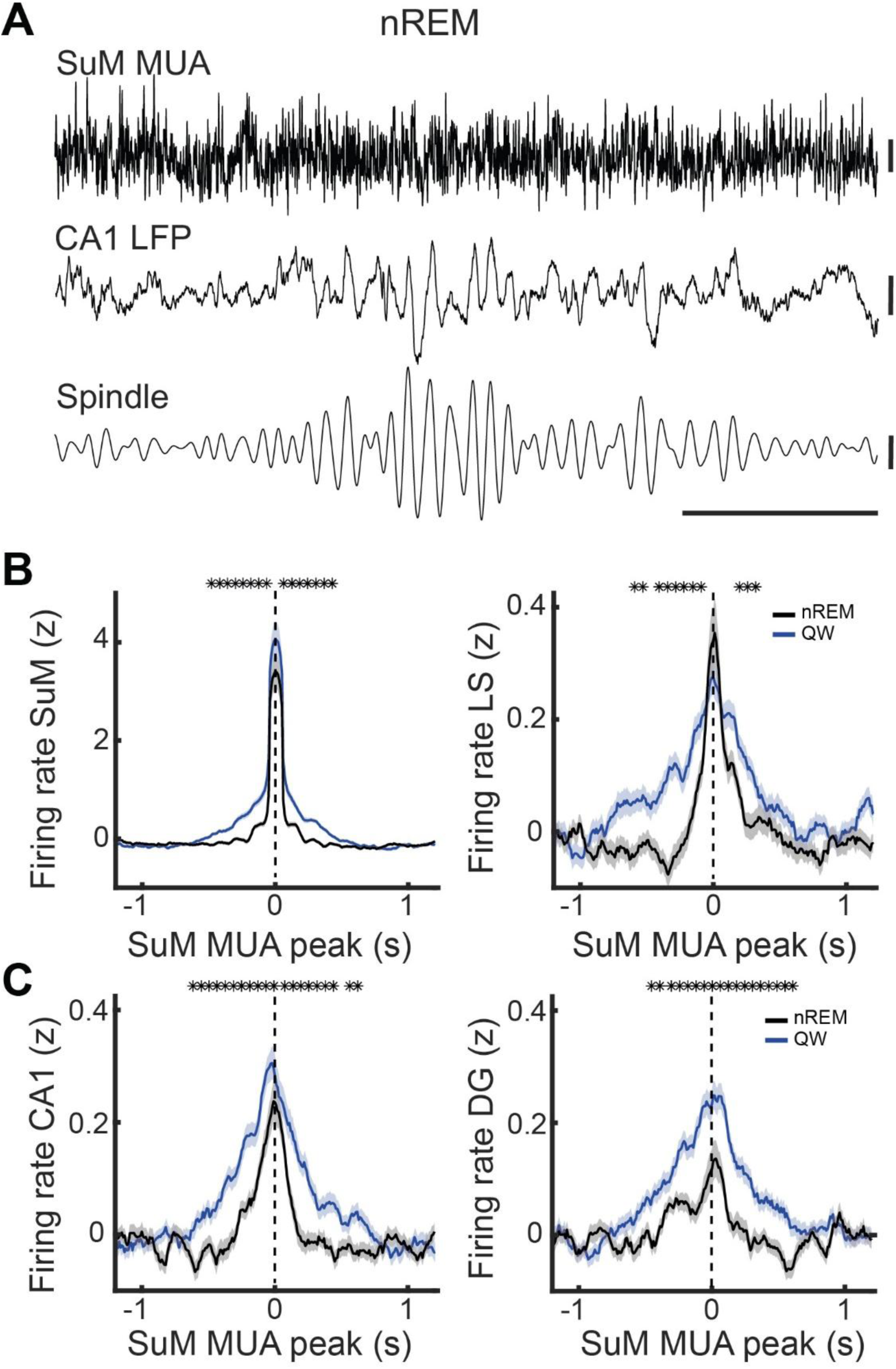
SuM burst–associated neuronal responses across hippocampal and subcortical populations during quiescent states. **A,** Example traces from the same session showing supramammillary nucleus (SuM) multi-unit activity (MUA), dorsal CA1 local field potential (LFP), and an example spindle from the CA1 LFP channel. Vertical bars depict amplitude scale; horizontal bar shows time scale. **B–C,** Peri-event firing-rate histograms (mean ± SEM; z-score) aligned to SuM MUA peaks (t = 0; dashed line), comparing non-REM sleep (nREM, black) with quiet wake (QW, orange). (B) SuM (left) and LS (right) show clear increases centered on SuM bursts, with larger responses in QW than in nREM around the peak response (centered at t = 0). (C) CA1 (left) and DG (right) also exhibit slower, modest activations time-locked to SuM bursts that are enhanced in QW relative to nREM. Asterisks above traces mark time bins significantly different between distributions (*p* < 0.05; FDR-corrected). QW, quiet wake; nREM, non-REM sleep; MUA, multiunit activity; SuM, supramammillary nucleus; LS, lateral septum; DG, dentate gyrus.

We next examined hippocampal–subcortical interactions during theta-enriched activated states, comparing REM sleep and active wakefulness (AW) (**Fig. 5A**). Aligning high-discharge SuM epochs to hippocampal activity revealed state-dependent effects, with SuM bursts producing rapid inhibition in CA1d selectively during wake (REM, 0.0 ± 0.04 z; AW, -0.22 ± 0.04 z; Wilcoxon signed-rank test, FDR-corrected, p < 1.8x10^-3^; **Fig. 5B**), whereas dorsal DG responses were similarly theta-modulated across brain-states (REM, 0.0 ± 0.04 z; AW, 0.0 ± 0.05 z; Wilcoxon signed-rank, FDR-corrected, p > 0.05; **Fig. 5B**). Next, to quantify neuronal frequency coupling, we computed the spike–field coherence of single units relative to the CA1 field potential. Neurons across regions showed a strong preference for the hippocampal theta band (**Fig. 5C**), and subcortical units (SuM and LS) exhibited stronger theta coupling in REM than in AW (**Fig. 5D**). Phase-locking analyses further revealed state-specific phase preferences and consequently sequential timing. Indeed, during wake, CA1 theta organized a descending-phase sequence, with DG firing first (mean angle = 111.6°, Rayleigh test, p = 1.3x10^-3^), followed by LS (mean angle = 146.2°, Rayleigh test, p = 9x10^-4^), and finally CA1 discharging near the trough (mean angle = 190.8°, Rayleigh test, p = 0.045; **Fig. 5E**). In such brain-state, SuM units were not significantly theta-modulated (mean angle = 326.2°, Rayleigh test, p = 0.12). Conversely, in REM, the pattern inverted and only SuM neurons were significantly phase-locked to CA1d theta (mean angle = 30.5°, Rayleigh test, p = 5.2 × 10⁻⁶), with preferred firing near the theta cycle peak (**Fig. 5E**). Together, these data show that theta-band coordination is reconfigured across activated states, with wakefulness favoring hippocampal firing around the trough of theta cycles and SuM bursts transiently inhibiting CA1, whereas REM selectively engages SuM coupling at the theta peak.

**Figure 5.**
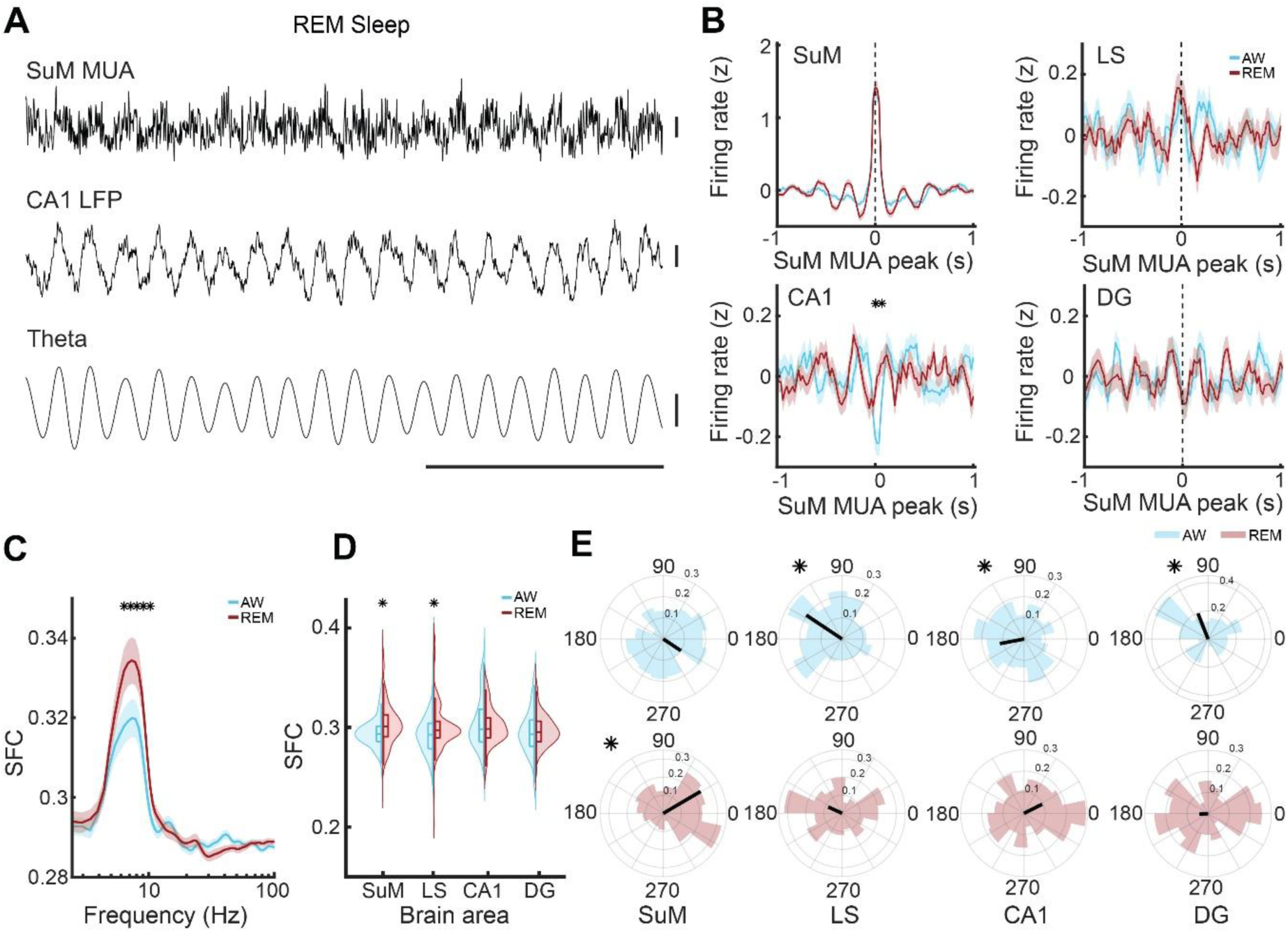
SuM burst–associated neuronal responses across hippocampal and subcortical populations during activated states. **A,** Example traces from the same session showing supramammillary (SuM) multi-unit activity (MUA), dorsal CA1 LFP, and the CA1 band-pass filtered theta reference. Scale bars at right (amplitude) and bottom (time). **B,** Peri-event firing-rate histograms (mean ± SEM; z-score) aligned to SuM MUA peaks (t = 0; dashed line) during active wake (QW, blue) and REM sleep (red). SuM exhibits a brief local peak (t = 0) in both states, and strong theta-modulation during REM; LS, CA1, and DG show smaller modulations. A transient suppression in CA1 around SuM peaks during AW is denoted (*). **C,** Spike–LFP coherence spectra (spikes referenced to the CA1 LFP) averaged across SuM units show stronger theta-band coherence (6–10 Hz) in REM than AW (asterisks indicate significant state differences, FDR-corrected). **D,** Theta-band spike–LFP coherence summarized by region (violin plots, box shows median and interquartile range). REM exceeds AW in SuM and LS (*p* < 0.05), with no robust state difference in CA1 or DG. **E,** Circular phase histograms of spike timing relative to theta (degrees) with mean resultant vectors (black arrows). LS, CA1, and DG exhibit significant theta phase-locking in AW; whereas SuM is not phase-modulated. Conversely, SuM shows significant theta phase-locking in REM, whereas LS, CA1, and DG display non-significant locking. AW, active wake; REM, rapid-eye-movement sleep; SuM, supramammillary nucleus; LS, lateral septum; DG, dentate gyrus. Asterisks mark FDR-corrected significant comparisons.

## DISCUSSION

Here we delineate how hippocampal–subcortical communication reorganizes across brain states, and the consequences of such dynamics for routing, timing, and gain control in extended hippocampal circuits (26, 28). Two organization motifs emerge from our data. First, during quiescent states (dominated by sharp-wave ripples) the hippocampus provides rapid top-down volleys to the associated subcortical nuclei, with CA1 ripples driving strong and DG dentate spikes more modest, but reliable, mostly state-invariant, excitation locally in CA1 and DG, and weaker excitation in LS and SuM. In the opposite direction, bottom-up streams from the SuM reach the hippocampus eliciting slow, state-dependent activation (10, 29, 30). Second, during activated states (theta waves-rich epochs), SuM burst epochs yield fast, brief inhibition of CA1 specifically during active wakefulness, yet theta phase-locking is selectively enhanced during REM sleep (25, 31, 32). These findings outline complementary effects, with fast ripple-locked top-down drive of SuM versus, SuM-mediated slow bottom-up modulation that reconfigures under theta oscillations.

Our results are consistent with the view that ripple episodes broadcast and entrain activity patterns well beyond the hippocampus that can bias downstream networks. Prior work has firmly linked ripples to systems-level coordination and selective engagement of large-scale networks, including nodes in the default mode network, consistent with extra-hippocampal responses that wax and wane with ongoing brain-state (5, 6, 33, 34). Our finding that hippocampal ripples evoke short-latency excitation in SuM and LS, but of smaller magnitude than locally in CA1 and DG, suggests effective but gain-limited propagation of the ripple volley into subcortical hubs. Such attenuated broadcast is compatible with the idea that ripples prioritize hippocampal–cortical consolidation while only sampling subcortical effectors (6, 28), thereby preventing indiscriminate state transitions during offline processing.

Another observation is the dissociation in SuM between state–dependent evoked potentials and largely state-invariant spike responses to ripples. In nREM, SuM field responses (i.e., event related potentials) are larger than in quiet wake, yet population peri-stimulus time histograms show remarkably similar spiking. This pattern implies that synaptic input to SuM scales with brain-state, but that local membrane potential, excitability mechanisms, or circuit architecture compress input–output gain, keeping spike output stable across quiescent states (35–37). Mechanistically, one possibility is that such a regime would allow the top-down transfer of cortical events via the hippocampus, without risking spurious bottom-up signalling of subcortical state transitions via SuM when the system is near sleep–wake transitions (28, 38, 39). Another option is that regardless of brain-state, subcortical nuclei like SuM, generate a consistent ascending output (38, 40). These observations also align with previous results of brain-wide analyses showing that ripple features covary with slower excitability fluctuations that differentially predict downstream responses (5, 6). On the other hand, dentate spikes presented an interesting contrast. Indeed, consistent with proposals that dentate spikes reorganize excitation–inhibition across the trisynaptic hippocampus circuit while exerting distinct network roles from ripples, we found that dentate spike-evoked responses were weaker locally and largely state-invariant, with little subcortical impact compared to ripples (13, 41, 42). Together, these observations support a division of labor among offline hippocampal events, with ripples furnishing potent, brief top-down broadcasts suited to rapid population coordination and replay; and dentate spikes providing subtler, possibly gating modulations that may tune entorhinal–hippocampal throughput without broadly driving subcortical systems (1).

Activated states dominated by theta oscillations revealed a distinct regime. Across REM sleep and active waking, spike–field metrics showed circuit-wide coordination by theta waves, but with significant state selectivity in SuM– hippocampus coupling. Classical single-unit studies established that SuM neurons fire rhythmically with hippocampal theta and, in many cells, near the theta cycle peak (43, 44). They further showed that SuM units often exhibit symmetric coherence with both CA1 and DG theta generators (45). Contrasting with such framework, our findings show that SuM phase-locking is robust in REM but weak or absent in active wake for the SuM neuronal population that we sampled. Moreover, SuM bursts in wake rapidly suppress CA1 spiking without equivalently suppressing DG, likely due to the selective activation of the GABAergic population SuM-innervated in the border between stratum radiatum and stratum lacunosum moleculare (46–48). Meanwhile, during REM sleep, SuM neurons are the only subcortical group showing significant theta phase-locking, with a preferred phase near the theta cycle peak. We interpret these results as evidence that SuM coupling to theta oscillations is dynamic, depending on brain-state and the sampled neuronal population. This may be an extension of recent evidence showing the differential tuning of mammillary neurons to hippocampal oscillations, which is strongly dependent on the precise location of the recorded neuronal populations within the medio-lateral axis of the mammillary bodies (6). According to our results, in REM sleep, SuM–hippocampus interactions may prioritize rhythmic coordination (i.e.; phase-locked discharge), whereas in wake, SuM can deliver rapid inhibitory control over CA1, while leaving DG timing largely governed by ongoing hippocampal theta.

Our observations integrate anatomical and physiological information. SuM provides dense glutamatergic/GABAergic input to DG, CA1, and CA2 and has collaterals reaching the septal complex (14–16). Moreover, classical work also established the capacity of SuM to pace or modulate hippocampal theta frequency and power (43, 49). Our state-dependent results align with a model in which SuM acts as a flexible coordinator. During REM, SuM neurons rhythmically entrain hippocampal timing; while in wake, brief SuM discharges impose fast, phase-specific inhibitory control in CA1. The observed asymmetry between CA1 and DG during wake-locked SuM bursts further suggests projection-defined selectivity or engagement of distinct microcircuits at these targets, possibilities consistent with heterogeneous SuM projections and interneuron recruitment previously reported (16). Our findings request specific mechanistic hypotheses. First, ripple-driven SuM activation engages synapses whose efficacy is state-dependent, given that ERPs are larger in nREM sleep; but seem to be embedded in a circuit with homeostatic gain control, yielding state-invariant spiking (50, 51). Second, during active wake, SuM bursts rapidly suppress CA1 via fast inhibitory pathways (e.g., hippocampal interneuron recruitment by direct innervation) while leaving DG relatively spared, thereby biasing information flow away from CA1 output . Third, during REM, SuM neurons preferentially lock to the theta cycle peak, positioning their discharge to influence CA1 timing, which spikes in the same theta phase, though not significantly, without strongly inhibiting DG and LS, that discharge at different phase; this configuration may align with known REM-linked SuM roles in arousal and theta pacing (38, 43).

In summary, hippocampal–subcortical communication is bidirectional but asymmetric in effects, time-scale, and brain-state dependence. Ripples confer rapid, high-gain top-down coordination with differential subcortical spiking: strong SuM and weak LS recruitment, while SuM provides slow, state-selective bottom-up modulation that reconfigures under theta oscillations, by suppressing CA1 spiking in active wake yet rhythmically coupling in REM. By integrating high-frequency oscillations controls with state-resolved spike–field analyses, these results refine canonical views of SuM–hippocampus coupling and situate ripple- and theta-based communication within a unified, state-dependent framework for extended hippocampal computation.

## METHODS

### Animals

Ten adult male Sprague–Dawley rats (postnatal day 40–60; 300–350 g) were obtained from the Center for Innovation on Biomedical Experimental Models (CIBEM, Pontificia Universidad Católica de Chile). Animals were housed at 22 ± 2 °C on a 12:12 h light/dark cycle (lights on 07:00), with food and water ad libitum. All procedures were approved by the Scientific Ethical Committee for the Care of Animals and the Environment (CEC-CAA, protocol 200610004). All methods complied with relevant guidelines and regulations and are reported in accordance with the ARRIVE guidelines.

### Habituation

Rats acclimated to the housing room for 2-3 days, then were handled daily by the experimenter for an additional 3–5 days. They were subsequently habituated to the recording environment by placement on a towel-covered drum in the recording room for 30–60 min per day over 1 week.

### Recording Implant Assembly

Electrode drives were designed in Autodesk Fusion and 3D-printed (Fusion3 F410). Each drive carried 16 tungsten, standard tapered-tip electrodes (MicroProbes, USA; 1, 3, or 5 MΩ) aimed at target regions defined from a rat stereotaxic atlas. Electrodes connected to EIB-18 interface boards (18 channels, including ground and reference). Ground and reference were attached to stainless-steel skull screws. The assembly was enclosed in copper mesh to reduce electrical noise.

### Stereotaxic Surgery

Anesthesia was induced with isoflurane (4%) and maintained at 1.5–2% in a stereotaxic frame. Body temperature was kept at 35– 37 °C with a homeothermic blanket; glucosaline (0.9% NaCl, 2.5% dextrose) was given hourly. After a midline scalp incision, four 1 mm craniotomies were made in the right hemisphere to reach the target areas at: CA1d: −4.2 AP, +3.6 ML, −2.9 DV; LS: 0.0 AP, 1.2 ML, −7.2 DV; DG: −4.2 AP, +3.0 ML, −3.8 DV; SuM: +2.5 AP, 1.5 ML, 9.3 DV (mm). Two additional anterior craniotomies housed ground and reference screws; four more (two contralateral parietal, one ipsilateral parietal, one posterior to lambda) anchored support screws. After durotomy, electrodes were lowered with a 10 degree angle away from the midline, craniotomies sealed with silicone elastomer or wax, and the drive secured with dental acrylic. Post-operative care included daily enrofloxacin (10 mg/kg, s.c.) and meloxicam (1 mg/kg, s.c.) for three days. Recordings began 7 days after surgery.

### Electrophysiological Recordings

Recordings were performed during the light phase for up to 10 consecutive days inside a Faraday cage. Rats sat on a towel-covered platform for 60–90-min sessions. The EIB-18 connected via a 16-channel Intan headstage to an Intan RHD amplifier; one electrode served as reference. Video (time-locked to the amplifier clock) aided brain-state scoring. Signals were sampled at 20 kHz using RHX (Intan Technologies) and converted to MATLAB with the LAN toolbox for analysis.

### Histology

At the end of experiments, rats were briefly anesthetized with isoflurane and small electrolytic lesions (5 µA, 10 s) marked electrode tips. After 48 h, animals were deeply anesthetized (ketamine 300 mg/kg + xylazine 30 mg/kg, i.p.) and perfused transcardially with 0.9% saline followed by 4% paraformaldehyde. Brains were post-fixed overnight, transferred to PBS-azide, sectioned coronally (vibratome; World Precision Instruments), Nissl-stained, and inspected (Nikon Eclipse CI-L) to verify stereotaxic placements.

### Brain-state Identification

Brain-states (quiet and active wakefulness, nREM and REM sleep) were scored in 10-s epochs using CA1 LFP (raw trace and Fourier spectrogram) and synchronized video (52). Quiet wakefulness featured mixed-frequency activity (>20 Hz) with minimal movement; active wakefulness was evidenced by conspicuous theta waves (5–10 Hz) and noticeable movement patterns; nREM displayed continuous low-frequency activity (0.1–4 Hz) without movement; REM showed robust theta oscillations (5–10 Hz) with muscle atonia. Custom MATLAB scripts implemented scoring following published methods.

### Oscillation Detection

Ripple episodes were detected by CA1 LFP band-pass filtering (100–250 Hz; zero-phase, non-causal, 0.5 Hz roll-off), rectified, and low-pass filtered at 20 Hz (4th-order Butterworth). After z-scoring, events crossing 3.5 SD and lasting ≥50 ms were detected and visually confirmed. Onset/offset were defined at 1 SD crossings; a 50 ms refractory prevented duplicates. Spectral features (frequency, amplitude, duration) were estimated with 7-cycle Morlet wavelets. Theta oscillations were detected by calculating the continuous ratio between the envelopes of theta (5–10 Hz) and delta (1–4 Hz) frequency bands filtered from the hippocampus LFP, and calculated by the Hilbert transform. A ratio of 1.5 SD or higher, during at least 2 s defined epochs of theta oscillations. Dentate spikes were detected by DG LFP band-pass filtering (100 and 250 Hz, 4th order Butterworth filter) and z-scored. High-frequency events based on amplitude, with 5–7 SD threshold where selected and visually confirmed.

### Time-frequency Analysis

The LFP from dorsal CA1 was considered as the time-frame reference for the spike-timing of recorded units. To determine the phase relationship between single-cell activity and theta cycles, the local field potential during theta episodes was filtered between 4 and 8 Hz, and the troughs of the theta oscillations were detected in the filtered signals.

### Cross-correlation Analysis

Activity of recorded units and LFP events (ripples, dentate spikes, SuM MUA) was cross-correlated by applying the “sliding-sweeps” algorithm (53). A time window of ± 1 s was defined with the 0 point assigned to the start time of a ripple. The timestamps of the unit spikes within the time window were considered as a template and were represented by a vector of spikes relative to t = 0 s, with a time bin of 50 ms and normalized to the total number of spikes. Thus, the central bin of the vector contained the ratio between the number of spikes elicited between ± 25 ms and the total number of spikes within the template. Next, the window was shifted to successive ripples throughout the recording session, and an array of recurrences of templates was obtained. Both unit timestamps and start times of LFP events were shuffled by randomized exchange of the original inter-event intervals (54) and the cross-correlation procedure was performed on the pseudo-random sequence. The statistical significance of the observed repetition of spike sequences was assessed by comparing, bin to bin, the original sequence with the shuffled sequence. An original correlation sequence that presented a statistical distribution different from 100 simulated shufflings was considered as statistically significant, with p < 0.01 probability, instead of a chance occurrence. For every recording with a significant correlation, the average of the simulated shuffling was subtracted from the average of the correlation curve and a representative cross-correlogram was obtained. To reveal repeating event correlations through the population, all representative cross-correlogram curves were pooled together and the statistical significance of a non-zero observed value was computed.

### Spike Sorting

Wideband data were band-pass filtered (600–5000 Hz) to extract spikes (negative or positive threshold crossings). Units were sorted with Kilosort2 (manual curation) using principal components. Putative pyramidal cells and interneurons were distinguished by waveform features and mean firing rate. Clusters were labeled single-unit if the refractory period was ≥1.5 ms; when isolation was ambiguous, clusters were conservatively classified as multiunit activity (MUA). For SuM MUA analysis, we selected LFP recordings from two closely located channels within SuM based on both anatomical location and LFP waveform characteristics. LFP signals were substrated (i.e., bipolar derivation), rectified (absolute value) and z-scored. Peaks were identified using the MATLAB (MathWorks) function *findpeaks* and peaks-amplitudes within the 0 – 4 SD range were extracted. These candidate MUA events were then temporally correlated with units previously sorted in SuM. The correlation analysis was used to determine the optimal detection threshold for MUA (Fig. S3).

### Statistical Analysis

Analyses followed standard parametric/non-parametric choices. One-way ANOVA tested single-factor group differences when normality and homoscedasticity held; otherwise, Kruskal–Wallis was used. Repeated-measures ANOVA was applied to repeated factors with normal residuals; paired t-tests compared two repeated levels, replaced by Wilcoxon signed-rank if non-normal; ≥3 repeated, non-normal levels used Friedman tests. Mauchly’s test assessed sphericity; Greenhouse–Geisser corrections were applied as needed. Post-hoc tests following ANOVA were Bonferroni-adjusted. Pearson’s r quantified linear relations for normal variables; Spearman’s ρ otherwise. Two-tailed α = 0.05 defined significance. All statistics were performed in MATLAB (MathWorks).

### Data Availability

Datasets are available from the corresponding author upon reasonable request.

## Acknowledgments

AI-assisted tools (ChatGPT) were used solely to improve clarity, grammar, flow, and to check statistical framing. All AI-assisted text was reviewed, edited, and validated by the authors. No core research tasks (experimental design, data acquisition, primary analyses, interpretation, or conclusions) relied on AI. The authors are fully responsible for the content.

## Author Contributions

PF designed the research and wrote the manuscript; NE performed experiments and analyses. GL, MC, AA, and AL-V performed experiments. The authors declare no competing interests.

---

**Figure S1.**
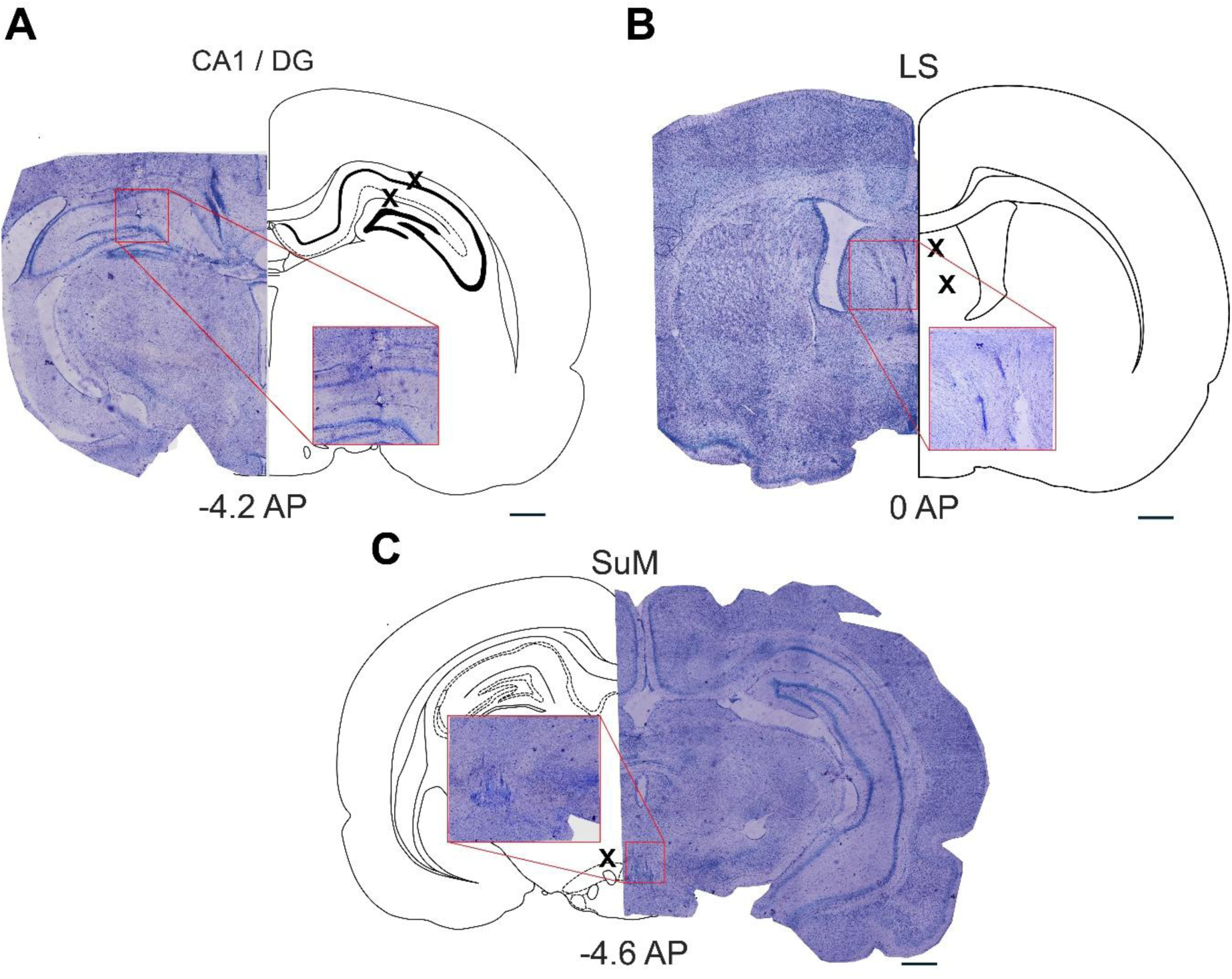
Histological verification of recording sites in hippocampus and subcortical targets. Coronal Nissl-stained sections from a representative animal confirming electrode placements in (A) dorsal hippocampus (CA1d and dentate gyrus, DG), (B) lateral septum (LS), and (C) supramammillary nucleus (SuM). For each target, a photomicrograph is shown alongside the corresponding atlas outline, with “×” symbols marking the estimated final recording sites. Insets show higher-magnification views of the electrode track/tip lesion in the target structure. Numbers indicate the antero-posterior (AP) coordinates relative to bregma printed on each panel (A: −4.2 AP; B: 0 AP; C: −4.6 AP). These reconstructions verify that hippocampal penetrations terminated within CA1d and DG, septal penetrations within LS, and hypothalamic penetrations within SuM. Scale bars shown. Abbreviations: CA1d, dorsal cornu ammonis 1; DG, dentate gyrus; LS, lateral septum; SuM, supramammillary nucleus; AP, anteroposterior.

**Figure S2.**
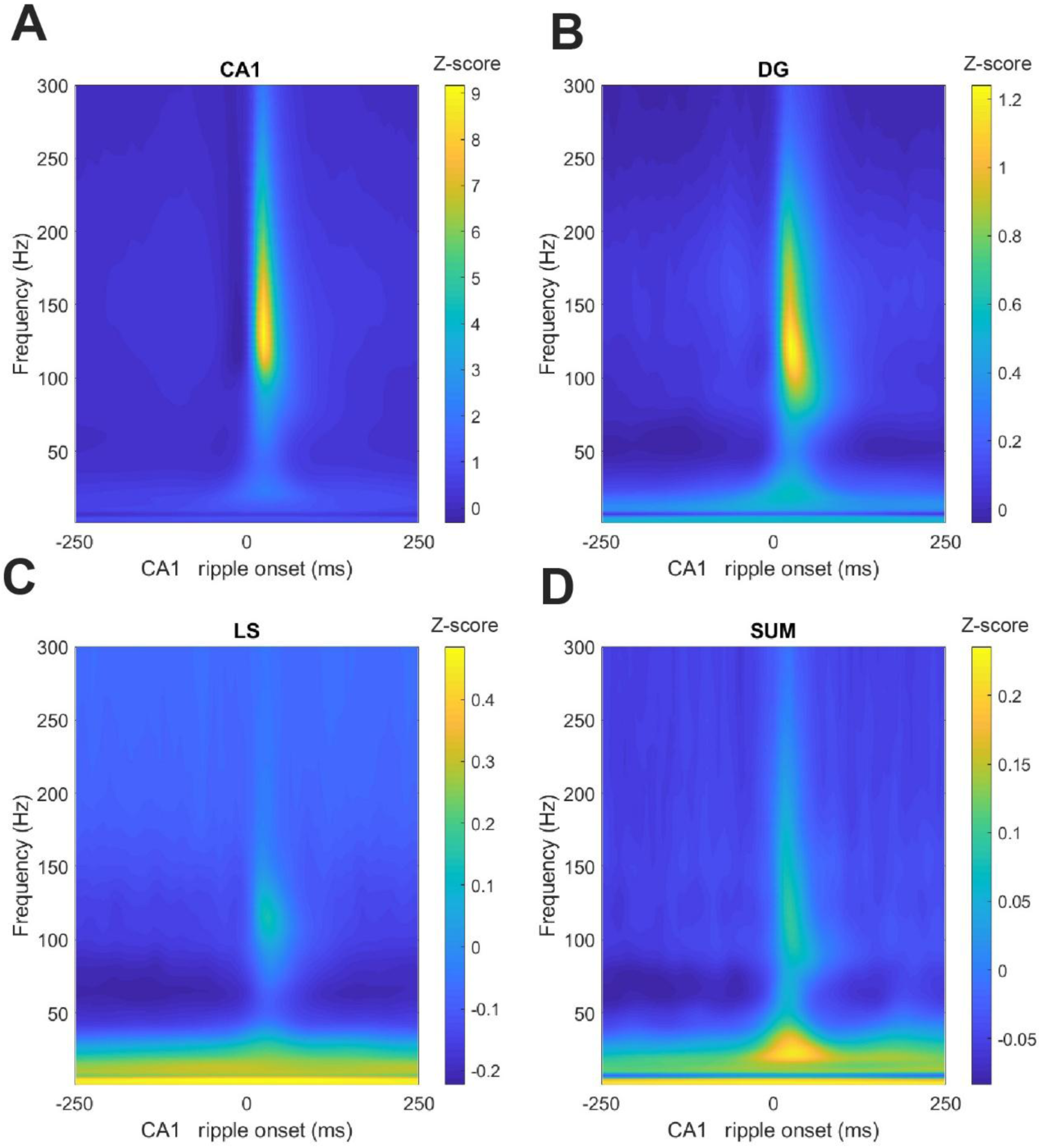
Ripple-triggered time–frequency structure across hippocampus and subcortical targets. Mean LFP spectrograms (wavelet time–frequency power; z-score relative to a pre-event baseline) aligned to CA1d ripple onset (t = 0; x-axis, −250 to +250 ms) for CA1d, dentate gyrus (DG), lateral septum (LS), and supramammillary nucleus (SuM). The y-axis spans 1– 300 Hz; separate colorbars (right of each panel) indicate the z-score range used for that region. CA1d exhibits the canonical narrow-band ripple increase (∼120– 180 Hz) centered at ripple onset, with a faint low-frequency (<30 Hz) component. DG shows a similar but smaller ripple-band increase. In contrast, LS and SuM display attenuated high-frequency power with only modest modulation (LS shows a weak 100–120 Hz bump; SuM is dominated by low-frequency evoked activity). Spectrograms are averages across all detected CA1d ripples and across sessions/animals. These profiles illustrate the strong local high-frequency expression in hippocampus and weaker, lower-frequency signatures in subcortical targets during ripple events. Abbreviations: CA1d, dorsal CA1; DG, dentate gyrus; LS, lateral septum; SuM, supramammillary nucleus.

**Figure S3.**
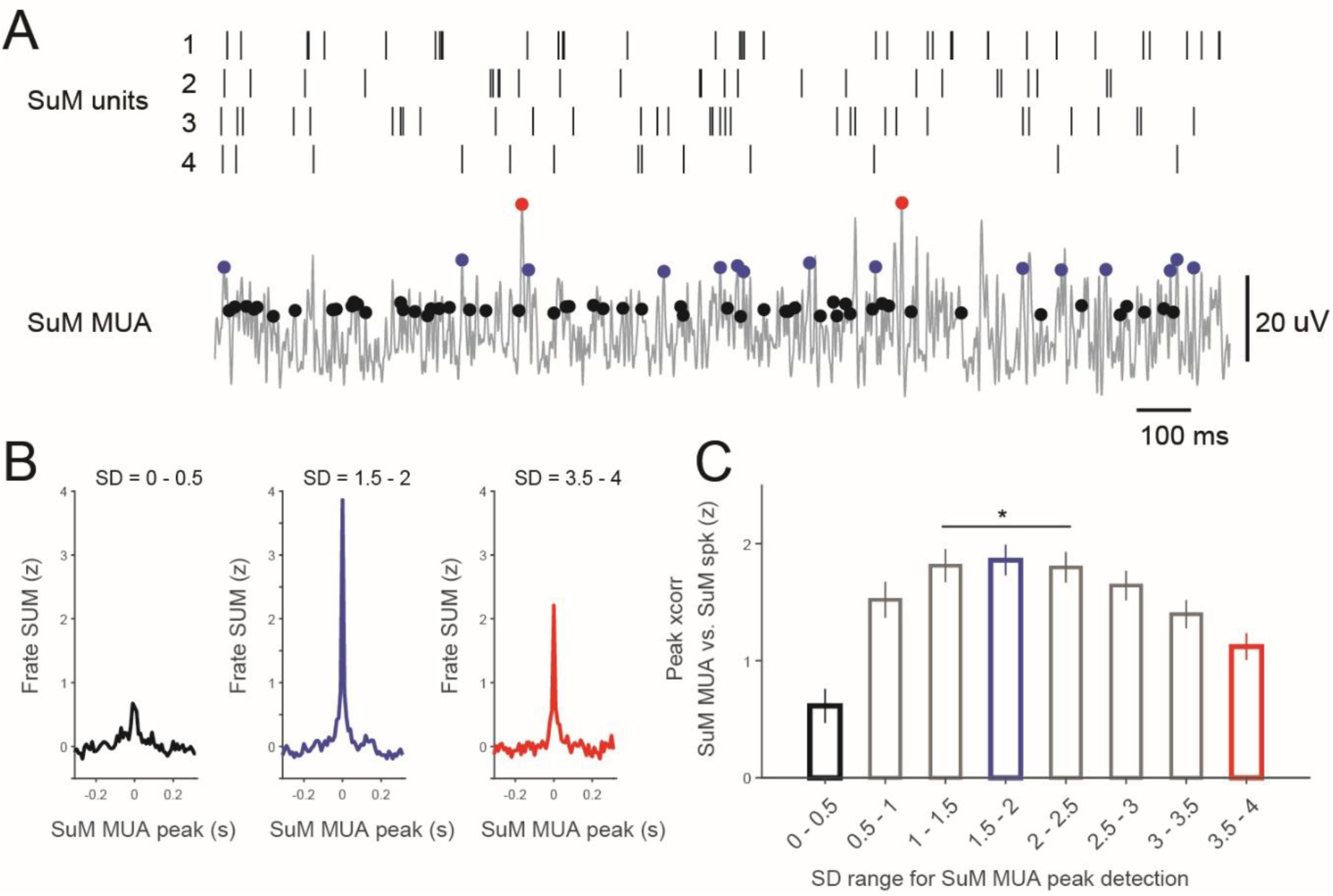
Selection of supramammillary (SuM) burst events from multi-unit activity (MUA). A, Example SuM recording illustrating the relation between simultaneously isolated SuM single units (rasters for four neurons, top) and the SuM MUA trace (bottom, gray). Colored dots mark peaks detected with three z-score threshold ranges: black = low (0–0.5 SD), blue = intermediate (1.5–2 SD), red = high (3.5–4 SD). Note that very low thresholds admit many small, noisy deflections, whereas very high thresholds capture only rare, large peaks. Scale bars: 20 µV, 100 ms. **B,** SuM peri-event firing rate (mean ± SEM; z-score) aligned to MUA peaks detected with the three thresholds shown in (A). Intermediate thresholds (1.5–2.5 SD) yield a sharp, symmetric burst centered at t = 0, whereas low thresholds (0–0.5 SD; left) give a broader, lower-contrast profile and high thresholds (3.5–4 SD; right) reduce event counts and attenuate the average. **C,** Peak cross-correlation (mean ± SEM; z-score) between the SuM MUA and the summed SuM single-unit spike train as a function of the MUA-peak threshold (binned in 0.5-SD steps). The correspondence between MUA peaks and local spiking maximizes at ∼1.5–2 SD and declines for very low or very high thresholds (*p* < 0.05; see Methods for statistics). These controls motivated using an intermediate z-score threshold to define SuM “burst” events in the main analyses.

**Table S1.** Histological identification of recorded brain regions per animal. ‘+’ indicates successful electrode placement in the region and inclusion of its data for analysis; whereas ‘–’ depicts failed electrode placement.

| Animal ID | CA1d | DG | LS | SUM |
| --- | --- | --- | --- | --- |
| AL07 | + | + | + | + |
| AL10 | + | + | + | + |
| MC07 | + | - | + | + |
| MC09 | + | + | + | + |
| MC14 | + | + | + | + |
| MC15 | + | + | + | + |
| MC16 | + | + | + | + |
| MC17 | + | - | + | + |
| MC18 | + | + | + | + |
| MC19 | + | + | + | + |

**Table S2.** Duration and relative proportion of brain states across animals during sleep recordings. Cumulative recording time (in minutes) per animal, together with the absolute (min) and relative (%) durations of nREM sleep, REM sleep, and quiet wakefulness (QW). Data represents totals across all recorded sessions. nREM mean and relative times are highlighted in bold.

| **Animal** | **QW**<br>**(min)** | **NREM**<br>**(min)** | **REM**<br>**(min)** | **QW**<br>**(%)** | **NREM**<br>**(%)** | **REM**<br>**(%)** |
| --- | --- | --- | --- | --- | --- | --- |
| **AL07** | 321.5 | 120.25 | 14.08 | 70.53 | 26.38 | 3.08 |
| **AL10** | 360.8 | 75.8 | 17.1 | 79.5 | 16.71 | 3.7 |
| **MC07** | 34 | 61.8 | 5.5 | 33.5 | 61.02 | 5.4 |
| **MC09** | 564.5 | 167.5 | 25.5 | 74.5 | 22.1 | 3.3 |
| **MC14** | 228.5 | 177.1 | 52.5 | 49.8 | 38.6 | 11.4 |
| **MC15** | 242.8 | 259.8 | 66.8 | 42.6 | 45.6 | 11.7 |
| **MC16** | 175.6 | 246 | 49.3 | 37.2 | 52.2 | 10.4 |
| **MC17** | 130.3 | 79 | 18.5 | 57.2 | 34.6 | 8.12 |
| **MC18** | 280.5 | 274.5 | 77 | 44.3 | 43.4 | 12.1 |
| **MC19** | 167.8 | 255.1 | 27.6 | 37.2 | 56.6 | 6.1 |
| **Mean** | 250.6 | 171.7 | 35.4 | 52.6 | 39.7 | 7.5 |
| **SD** | 145.6 | 83.9 | 24.3 | 16.8 | 14.8 | 3.6 |

**Table S3.** Total recording sessions and unitary activity recorded per animal and brain region. SUA stands for single unit activity. MUA stands for multi-unit activity. Units refers to the sum of all recorded SUAs and MUAs per animal.

| **# units (SU & MU)** |  |  |  |  |  |
| --- | --- | --- | --- | --- | --- |
|  | **# sessions** | **CA1d** | **DG** | **LS** | **SUM** |
| **AL07** | 6 | 10 | 16 | 17 | 15 |
| **AL10** | 3 | 0 | 16 | 0 | 15 |
| **MC07** | 1 | 1 | 4 | 0 | 13 |
| **MC09** | 6 | 18 | 15 | 16 | 26 |
| **MC14** | 7 | 40 | 31 | 19 | 29 |
| **MC15** | 9 | 40 | 43 | 21 | 23 |
| **MC16** | 9 | 44 | 39 | 30 | 35 |
| **MC17** | 4 | 19 | 9 | 14 | 3 |
| **MC18** | 8 | 30 | 0 | 33 | 29 |
| **MC19** | 6 | 35 | 6 | 23 | 35 |
| **Total** | **59** | **237** | **179** | **173** | **223** |

